# The threshold for intracranial self-stimulation does not increase in rats exposed to chronic unpredictable stress - a systematic review and meta–analysis

**DOI:** 10.1101/2024.01.15.575675

**Authors:** Jenny P. Berrio, Jenny Wilzopolski, Katharina Hohlbaum, Otto Kalliokoski

## Abstract

The chronic unpredictable stress model is a laboratory rodent model of stress-induced anhedonia. The sucrose preference test, often used to validate it, suffers from being unreliable. Intracranial self-stimulation offers an alternative and is often cited as supporting evidence of the validity of the model. Our aim was to assess whether an increased self-stimulation threshold is found after stress and if such a change correlates with decreases in sweet consumption. We searched PubMed, Embase, and Web of Science for studies in rats exposed to chronic unpredictable stress that employed intracranial self-stimulation. Thresholds, for stressed and control animals, in 23 experiments (11 studies) were pooled. Over 50% of the data was contributed by one research group, so a three-level meta-analytical random effects model was fit to account for methodological differences between different networks of researchers. After this adjustment, we did not find that the self-stimulation thresholds were increased in stressed rats. Pioneering experiments with positive results failed to be replicated by others, although no specific factor could be pointed to as a likely explanation. What is more, the available evidence suggests a lack of connection between sweet preference and self-stimulation, although this relationship has been seldom investigated. Methods known to mitigate biases were frequently absent, as was a transparent report of crucial study details. Our findings challenge the claim made in support of the validity of the model. Further efforts would be well-invested in assessing how reliably other tests of anhedonia have found the effects of the chronic unpredictable stress model.

## Introduction

The term anhedonia was described over a century ago by Théodule Ribot^1^ as “the inability to experience pleasure”. Even though recent advancements in the understanding of the biology behind the experience of pleasure have highlighted the need to expand this definition to reflect its complexity^2^, this broad conceptual frame is still largely used in clinical and preclinical research^3^. Anhedonia (defined as “markedly diminished interest or pleasure in all, or almost all, activities”) is listed in the Diagnostic and Statistical Manual of Mental Disorders (DSM-5)^4^ as one of two major symptoms needed to diagnose a Major Depression episode^4^. A patient is considered to be suffering from a depressive episode when five of the characteristic symptoms of depression are present for a minimum of two weeks, one of which must be either a depressed mood or anhedonia. Since depressed mood is a subjective experience, difficult to assess in non-human species, many models aiming to mimic depression have focused on reproducing anhedonia in the laboratory.

Animal models based on chronic exposure to stress are widely used for modeling depression. The idea is based on an accepted link between the repeated exposure to adverse life events and the risk of developing depression in humans^5^. In particular, the chronic unpredictable stress (CUS) model, one of the most common models of depression in laboratory rodents^6^, makes use of environmental stressors (*e.g.*, social crowding or isolation, food and water deprivation, wet bedding, light changes) to create adverse situations^7^. These stressing stimuli are delivered constantly, albeit unpredictably, over several weeks, and sometimes even months. This model was developed following the observation that rats exposed to stressors decreased their consumption of a sweet solution^8^. This effect was interpreted at the time to mean that the capacity of the animals to enjoy the solution had decreased; they had become anhedonic. Thus, rodents exposed to such protocols are expected to decrease their consumption of a mildly sweetened solution in the so called sucrose preference test (SPT)^6^. The assumption is that the preference for the sweet solution corresponds to the pleasure derived from consuming it. Therefore, when stressed animals consume less solution compared to unstressed conspecifics, it is interpreted as a lower capacity for experiencing pleasure. This is the test most commonly used to assess anhedonia in rodents^3^ and the one used to demonstrate that the model reliably induces anhedonia^9,10^. However, the reliability of this method has been questioned^6,9,11,12^. In fact, our recent systematic review of studies of laboratory rats exposed to CUS found high variability in the results of the test across different experiments. Moreover, we found methodological limitations that undermine our confidence in the test as a reliable way for accurately assessing or measuring anhedonia^10^.

Intracranial self-stimulation (ICSS) is an alternative method for assessing anhedonia. The extent of sweet consumption and self-stimulation are both considered measures of how much a reward is liked, only the type of reward is different. ICSS is an operant task in which rats learn to perform an action in order to stimulate their own brain^13^. More specifically, they learn that by manipulating an object such as a lever or a wheel, they can send an electrical stimulus via implanted electrodes. These electrodes are positioned in brain areas that belong to the neuronal network that processes reward. The electrical stimulation makes the neurons in these centers fire, evoking a rewarding experience and rats will self-stimulate (*e.g.,* continue lever pressing) to get more. Whether true pleasure is experienced is still up for debate. Berridge and collaborators have made the case that electrodes enhance the motivation to pursue the reward (wanting the simulation) without necessarily producing feelings of pleasure^14^. Nevertheless, humans and rodents alike will respond avidly to such stimulation^13,15,16^. Self–stimulation thresholds are a quantitative measure of stimulation efficacy and thus serve as a means to understanding how different factors affect the brain’s pleasure system^17^. If the threshold is low, it means the brain’s reward circuitry is very sensitive, and even a weak stimulation is perceived as “pleasurable”. If the threshold is high, it means the brain’s reward circuitry is not as sensitive and the animal needs to be exposed to a stronger stimulation (*e.g.* a higher frequency or intensity) in order to start perceiving it as rewarding. Shifts in the self-stimulation threshold in response to a particular intervention or treatment are likewise informative. In particular, an elevation in the threshold, say in response to chronic stress, indicates stress has reward-attenuating effects (interpreted as stress-induced anhedonia). This effect has been reported for the CUS model and it is often cited in support of the validity of the model in reproducing anhedonia amidst concerns regarding the challenges in replicating the results of the SPT. The supporting argument emphasizes that chronically stressed rats, aside from exhibiting changes in their sweet consumption, show abnormal behaviors in other tests of anhedonia. In one of his latest publications, Paul Willner, who co-created the model, explains that the fact that CUS causes a general decrease in reward sensitivity is uncontroversial because the model, among other things, has been shown to increase the threshold for self-stimulation^6^.

There are several advantages of the intracranial self-stimulation procedure over other methods that use natural rewards such as food. First, because the procedure involves direct stimulation of the brain’s reward centers, it bypasses some of the steps related to the processing of sensory inputs^16,17^. In the case of sweet rewards, for example, the animal’s taste buds or brain might process the message in ways that may modify the response towards sweet food and confuse its interpretation^16^. For example, hunger and other metabolic cues can increase the hedonic value of food by regulating dopamine release in the dorsal striatum, a mechanism designed to prioritize energy seeking over taste quality^18^. By circumventing these steps using ICSS, we avoid confounding factors that are not related to the likeability of the reward. Second, the stimulation can be controlled to a finer level than with natural rewards. For example, different parameters of the stimulation can be individually changed to assess how this change affects self-stimulation behavior. Changes in frequency alter the pattern of firing of the neurons in the stimulated area while changes in current intensity recruit more neurons^17^. Additionally, since the electrodes can be implanted in different areas of the brain, it is possible to assess how different parts of the reward system are affected by the intervention^17^. Third, the response is believed to be unaffected by satiation or anxiety^19^. Rats, when given the choice, will engage in brain stimulation of reward centers even at the expense of self-starvation or while sustaining painful shocks^20^. Last but not least, ICSS thresholds are very stable in well-trained animals^16,19^. This is of great importance because it allows the use of within-subject designs in experiments and the evaluation of the effects of long exposures to interventions like CUS^16^.

Despite the strengths of ICSS and its role in supporting the validity of the model, to our knowledge, no systematic review of the effects of CUS on ICSS outcomes have been conducted to date. In the present study we aimed to assess whether chronically stressed rats exhibit an increased self-stimulation threshold and if such a change correlates with decreases in sweet consumption. Our focus was centered on rats, as this species was originally employed to develop the CUS model. Moreover, these studies in rats are often cited as evidence of the effect of the model on ICSS. We conducted a systematic review of studies in rats exposed to CUS that were assessed for intracranial self-stimulation. We hypothesized that: (1) stressed rats should show an increased ICSS threshold compared to their pre-stress values and in comparison to unstressed controls. (2) If this decreased sensitivity towards reward is also assessable through the SPT, then both variables should co-vary, and an increased threshold should correlate with a decreased sweet consumption. For this latter hypothesis, we were only interested in studies where a SPT was carried out in the same cohort of animals that were assessed for intracranial self-stimulation.

## Methods

### Registration and open access data

The protocol was registered in PROSPERO (CRD42023425916). The report adheres to the PRISMA guidelines for reporting^21^. The supplementary material provides in-depth information on the methods, along with links to a data repository. Deviations from the registered protocol are listed in detail in the supplementary materials.

### Search strategy

Relevant publications from 1987 (the year the model was created) to May 2023 were retrieved from three databases: PubMed, Embase and Web of Science. For all databases, three strings were created and combined. The combined search string looked for studies in (1) rats, (2) using CUS and (3) performing intracranial self-stimulation (ICSS). No other constraints were used.

### Screening Platform

The Covidence systematic review software (Veritas Health Innovation, Melbourne, Australia) was used for the screening process and data extraction^22^.

### Eligibility criteria

Following the removal of duplicates, each publication was screened by two independent reviewers in two phases. An initial title and abstract screening excluded studies that: (a) did not describe an original study in laboratory rats, (b) used other means to induce chronic stress and (c) did not use intracranial self-stimulation. In cases where there was disagreement between reviewers or incomplete information for exclusion, the study was passed on to the next phase. In the second phase, the full-texts of the remaining studies were screened. Studies that performed CUS in inbred and outbred weaned rats for at least two weeks and that used ICSS to assess anhedonia were included. A third reviewer, casting a deciding vote, solved discrepancies at this stage. To identify studies that might have been missed in our initial search, the reference lists of the included studies were checked. However, no additional studies were found this way.

### Study details and outcome data extraction

Supplementary table 2 presents the study details and outcomes extracted from each included study. Two reviewers independently performed the extraction, while a third reviewer resolved discrepancies. In cases where a study reported multiple experiments (different cohorts of animals) that met the inclusion criteria, each experiment was extracted separately. The stimulation thresholds before and at the end of CUS (mean, sample size, standard deviation or standard error) were extracted for animals exposed to stress. Likewise, if these data were reported for control (unstressed) animals, they were also extracted. Two additional outcomes were of interest for the present study. In the event of a study performing the SPT in the same cohort of animals in which intracranial self-stimulation was assessed, the sweet consumption/preference before and after stress were to be extracted. Furthermore, if the study included a correlation analysis between self-stimulation and sweet preference, the correlation coefficient was extracted as well. Whenever possible, information was collected from original data, tables, or text. Data presented in figures were extracted using a digitizing tool^23^. In these cases, the two values obtained by the reviewers were averaged and the mean value was used in the analyses. Supplementary table 3 outlines the considerations adhered to while conducting the extraction of outcome data according to different study methodologies and ways of reporting data. If a study had multiple control or stress groups and it was not possible to directly extract a “one-to-one” comparison, we followed the guidelines outlined in the Cochrane Handbook^24^ for combining multiple groups into one. Efforts were made to contact authors in case of missing data. When no response was received and other options for obtaining the data were exhausted, the study/experiment was excluded from the analysis if outcome data were incomplete.

### Risk of bias assessment

The methodological quality of the studies included in the study was assessed through a checklist adapted from the SYRCLE’s risk of bias tool^25^. Two reviewers evaluated each study independently focusing on nine items assessing selection, performance, detection, attrition and reporting bias (Supplementary table 4). Each item was evaluated by answering signaling questions that aimed to assess whether measures that mitigate each of these biases were employed in the studies. These questions were answered either with ‘Yes’ (indicating low risk), ‘No’ (indicating high risk) or ‘Unclear’. When there was disagreement, a third reviewer resolved the conflict. Each positive response was assigned one point, and the total points were tallied up for an overall risk of bias score. Since this assessment was conducted for each study (not at the experiment level), all experiments derived from a single study received the same score, and this score was used in the meta-analysis (see Data synthesis below). The traffic light plot was created using Robvis^26^.

### Data synthesis

The statistical analysis and the creation of the plots were performed in RStudio 2022.07.2.576^27^ with R version 4.2.1^28^. The full list of packages used is presented in the supplementary material.

A meta-analysis was considered possible when a minimum of four studies were included per outcome. We did not exclude studies based on a high risk of bias. Whenever the standard error was reported, it was converted to standard deviation for the meta-analysis. If the sample size in a study was reported as a range, the analysis was based on the lowest value in the range to have a conservative approach. We combined two stress groups in only one experiment. The effect of stress on intracranial self-stimulation was evaluated by comparing pre- and post-stress thresholds in stressed animals and by comparing post-stress thresholds in control and stressed animals. In this latter case, and in lieu of discarding experiments that only reported data for stressed animals, the pre-stress values were assumed to be the control (four experiments, details in the results section). For these cases, the group size was split in two to avoid double-counting^24^. Pooling and weighting of individual effect sizes (standardized mean differences, SMD - Hedges’ g) was done with a random-effects model using the restricted maximum likelihood (REML) for estimating between-study heterogeneity (referred to from hereon as the two-level meta-analysis). During the data extraction, we observed that several of the included studies were published listing the same first author. Moreover, we observed that several publications authored by different people used similar methods. Therefore, the list of authors of each included study was checked to define co-authorship networks based on collaboration. A network was formed when at least one author in a paper was listed as an author in another. Because some groups may be more inclined to find specific outcomes due to methodological factors or ways of analyzing experiments, effect size estimations might be distorted if this is not taken into account^29^. To control for the influence of the co-authorship network, we conducted a three-level meta-analysis using this variable as an additional nested level. To determine if this model better suited our data, the results of the three-level meta-analysis were compared with those obtained with the two-level meta-analysis using model fit statistics and a likelihood ratio test. For the three-level meta-analysis, different ICSS outcome measures were pooled and weighted (SMD) with a random effects model using the REML method for estimating τ^2^ at level 2 (within co-authorship network) and at level 3 (between co-authorship networks). The statistical measure for assessing heterogeneity was I^2^. Thus, two heterogeneity measures were calculated, each quantifying the percentage of total variation associated with either level 2 or 3 (I^2^ level 2, and I^2^ level 3, respectively).

### Exploring heterogeneity

At least four studies per subgroup were deemed necessary to conduct subgroup analyses. None of the categorical variables of interest (Supplementary table 2) had enough studies in the different subgroups to explore their influence on the results. The effect of continuous variables (length of stress and risk of bias score) was explored through multi-level meta-regressions.

### Sensitivity analyses and publication bias

To test the robustness of our findings two sensitivity analyses were conducted. In the first one, the analyses were re-run considering the sample size of those studies that reported the sample size in ranges, using the highest number instead. In the second, the data was re-analyzed after excluding experiments without a control group. The influence of small studies was explored through the visual inspection of a funnel plot and Egger’s test with the corrected standard error (SE) proposed by Pustejovsky and Rogers^30^.

### GRADE assessment

The confidence in the body of evidence was assessed using the GRADE approach^31^ by consensus among the reviewers. GRADE categorizes the certainty of evidence as high, moderate, low, or very low (Supplementary Table 5) based on the type of studies that form the evidence base and through the assessment of several factors related to quality and precision of the evidence. Factors such as risk of bias of the included studies, the degree of heterogeneity in effect sizes across experiments (inconsistency), the precision of the overall effect (imprecision), the indirectness of the evidence, and publication bias were assessed to determine the certainty of the evidence. The GRADE assessment was performed separately for the estimate obtained with the two-level meta-analysis and the three-level meta-analysis.

## Results

### Screening process

Figure 1 presents the PRISMA flow diagram detailing the selection process, the number and reasons for exclusions in each phase, and the number of studies included in our investigation. Out of the 195 non-duplicated records found through our search, 11 studies described 23 original experiments on laboratory rats exposed to CUS for at least 2 weeks and tested using intracranial self-stimulation.

**Figure 1.**
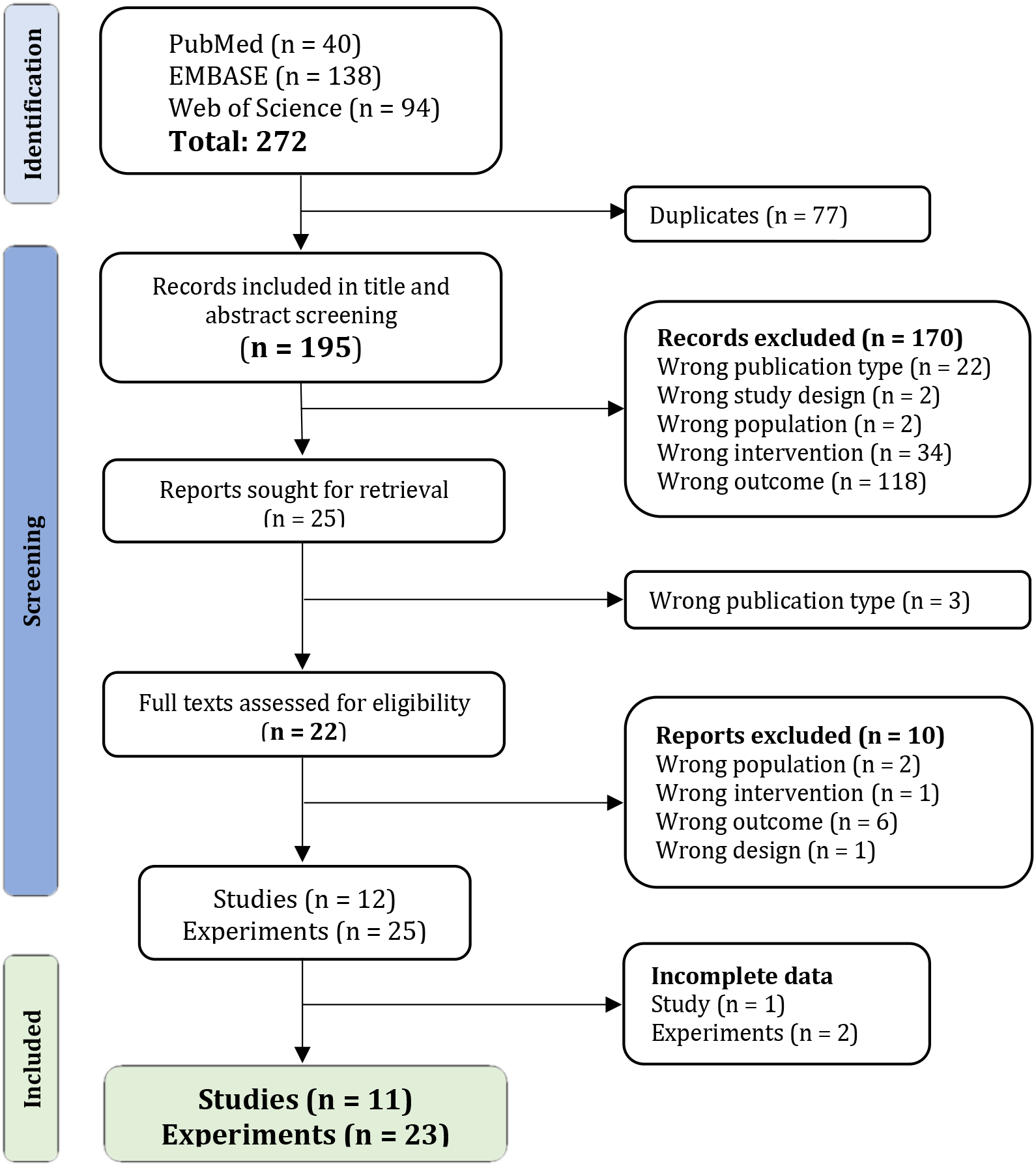
PRISMA flow diagram. Overview of the process for both the screening of titles and abstracts, as well as the comprehensive evaluation of full texts. The combined search across all databases produced 272 reports. After duplicates were removed, 195 reports were screened by title and abstract. Of those, only 25 reports passed on to the next phase. Twelve records met all the criteria following full-text review. One study^32^ did not have complete data, while another reported two experiments, one of which had incomplete data. After exhausting the options to acquire the missing information, these two experiments were excluded. A total of 11 studies and 23 unique experiments were included in the analysis. Adapted from the “PRISMA 2020 flow diagram for new systematic reviews”^21^.

### Characteristics of the included studies

#### Subject characteristics

All the studies included in the analysis were published between 1992 and 2006. Of the experiments, 52% (12 experiments from 6 studies) were published by the same first author and marked the initial contributions on the subject, spanning from 1992 to 1996^32–37^. Only four experiments used baseline measurements from the stressed group as the control condition (*within-subject design*), the other experiments employed unstressed rats as controls (*control vs exposed design*). Almost 70% of the experiments (16) were conducted using male Wistar rats, with the most substantial subset comprising Wistar (RoRo) rats (**Table 1**). Only four experiments employed female rats. Young adult rats were used in 83% of the experiments (19). The duration of the CUS was varied, ranging from 2.7 weeks to 9 weeks.

**Table 1.** Subject characteristics of the included experiments.

| Experiment | Co-authorship network | Design | Strain/Stock | Sex | Age | Weight (g) | CUS length (wks) |
| --- | --- | --- | --- | --- | --- | --- | --- |
| Baker 2006 C1 | Baker-Bielajew | Within-subject | Sprague Dawley | F | Adult 12 mos or less | 242-329 | 3 |
| C2 |  | Within-subject | Long Evans | F | Adult 12 mos or less | 242-329 | 3 |
| Bielajew 2003 C1 |  | Ctrl vs Exp | Sprague Dawley | F | Adult 12 mo or less | 220-300 | 6 |
| C2 |  | Ctrl vs Exp | Sprague Dawley | M | Adult 12 mos or less | 300-505 | 6 |
| C3 |  | Ctrl vs Exp | Long-Evans | F | Adult 12 mos or less | 220-300 | 6 |
| C4 |  | Ctrl vs Exp | Long-Evans | M | Adult 12 mos or less | 300-505 | 6 |
| Lin 2002 C2 | Lin |  | Wistar | M | 9 wks or less | 250-300 | 2.7 |
| Moreau 1992 | Moreau -Stout | Ctrl vs Exp | Wistar (RoRo) | M | Adult 12 mos or less | 350 | 2.7 |
| Moreau 1993 |  | Ctrl vs Exp | Wistar (RoRo) | M | Adult 12 mos or less | 350 | 2.7 |
| Moreau 1994b C1 |  | Ctrl vs Exp | Wistar (RoRo) | M | Adult 12 mos or less | 350 | 2.7 |
| C2 |  | Ctrl vs Exp | Wistar (RoRo) | M | Adult 12 mos or less | 350 | 2.7 |
| C3 |  | Ctrl vs Exp | Wistar (RoRo) | M | Adult 12 mos or less | 350 | 2.7 |
| Moreau 1994 C1 |  | Ctrl vs Exp | Wistar (RoRo) | M | Adult 12 mos or less | 350 | 2.7 |
| C2 |  | Ctrl vs Exp | Wistar (RoRo) | M | Adult 12 mos or less | 350 | 5.4 |
| Moreau 1995 |  | Within-subject | Wistar (RoRo) | M | Adult 12 mos or less | 350 | 4.7 |
| Moreau 1996 C1 |  | Ctrl vs Exp | Wistar (RoRo) | M | Adult 12 mos or less | 350 | 2.7 |
| C2 |  | Ctrl vs Exp | Wistar (RoRo) | M | Adult 12 mos or less | 350 | 2.7 |
| C3 |  | Ctrl vs Exp | Wistar (RoRo) | M | Adult 12 mos or less | 350 | 2.7 |
| C4 |  | Within-subject | Wistar (RoRo) | M | Adult 12 mos or less | 350 | 6 |
| Stout_2000 |  | Ctrl vs Exp | Wistar (RoRo) | M | Adult 12 mos or less | 350 | 2.7 |
| Nielsen 2000 C1 | Nielsen | Ctrl vs Exp | Wistar | M | 9 wks or less | 250-300 | 9 |
| C2 |  | Ctrl vs Exp | Wistar | M | 9 wks or less | 250-300 | 9 |
| C3 |  | Ctrl vs Exp | PVG | M | Unknown | 220-240 | 9 |

**Abbreviations:** wks, weeks; mos, months; Ctrl vs. Exp, control vs exposed, F, female; M, male; vs, versus “Cx” represent the number of experiments (comparisons) extracted from a single published study.

In our study, a co-authorship network was formed when authors had collaborated on one or more publications. Four networks were identified when the list of authors of the included studies was reviewed (Figure 2). The studies were grouped according to these networks: ‘Moreau-Stout’ comprising studies published by Moreau et al.^32–37^ and Stout et al.^38^ (seven studies with thirteen experiments), ‘Baker-Bielajew’ including studies published by Baker et al.^39^ and Bielajew et al.^40^ (two studies with six experiments), and ‘Lin’ and ‘Nielsen’, each consisting of one study (with one^41^ and three experiments^42^, respectively).

**Figure 2.**
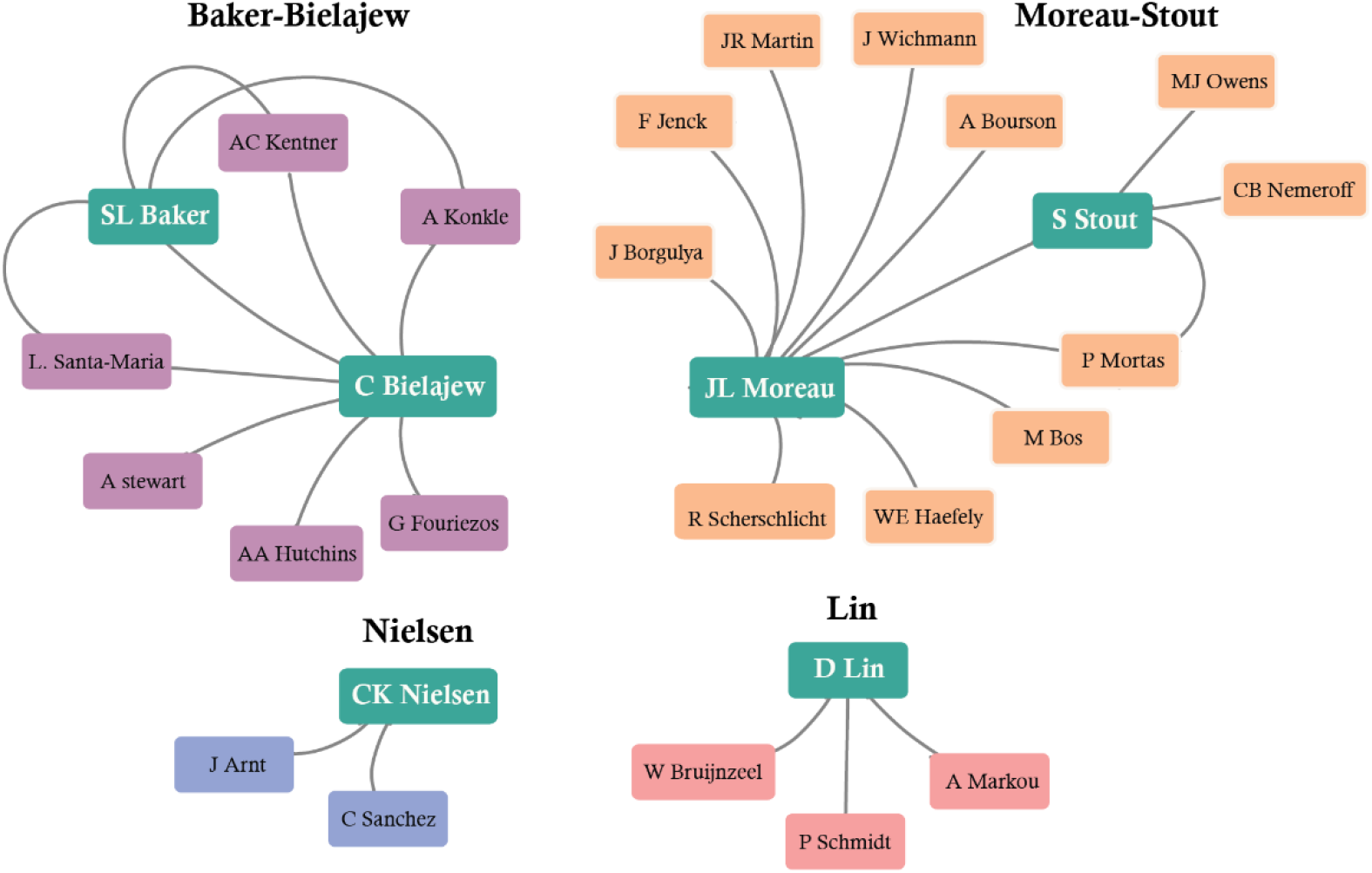
Co-authorship networks. The first authors of the included papers are shown in green boxes, the arrows connect these first authors to authors they collaborated with in one or more publications. These secondary authors are shown in boxes of different color according to the network they belong to. The ‘Moreau-Stout’ network is composed of the authors of 7 studies, ‘Baker-Bielajew’ includes the authors of 2 studies and ‘Lin’ and ‘Nielsen’ are composed of authors that collaborated on a single study.

#### ICSS methodology

**Table 2** summarizes the ICSS methodology employed in the experiments by presenting it by co-authorship network. Most networks, except one, chose the ventral tegmental area (VTA) as their target area for self-stimulation. Stereotaxic coordinates for this area varied among co-authorship networks and targeted different subregions. Nielsen’s coordinates match with what Paxinos and Watson^43^ identify as the parabrachial pigmented and the parainterfascicular nuclei; Baker-Bielajew’s coordinates match with the rostral part of the VTA, while Moreau-Stout’s coordinates match a more posterior subregion simply referred to as the VTA in the Paxinos and Watson’s atlas^43^. Three networks implanted bipolar electrodes that delivered either cathodal stimulation or stimulation of alternating polarity. The remaining network (Baker-Bielajew) implanted a monopolar electrode that delivered only cathodal stimulation. The object which the animal interacted with in order to trigger the stimulation (the manipulandum) was different between networks and included a lever to press, a wheel to rotate, and a hole for the animal to poke its snout through. In all studies, the electrical stimulation consisted of either a 300 or 500 millisecond train of pulses. The rate-frequency curve shift procedure was chosen by most networks for defining reward thresholds. During this procedure, different frequencies were tested in a one or two-minute trial. Thresholds were defined as either the M50 (the stimulation frequency at which the animal responds at a rate that is half their maximum response) or the frequency eliciting 15 nose-pokes per minute (Moreau-Stout). Lin and collaborators employed a discrete-trial procedure with a response window of 7.5 seconds to obtain the mean intensity at which the animal starts to respond positively to stimulation in the middle forebrain bundle. The measures reported were either the absolute frequency (Baker-Bielajew), or a measure of change post-stress: a percentage change from baseline (Moreau-Stout and Lin) or the average change (Nielsen). Overall, the Moreau-Stout and Nielsen networks had the most similar methodology, differing only in the threshold measure and the specific stereotaxic coordinates used to target the VTA. Nielsen and colleagues’ threshold was defined as the frequency supporting 50% of the maximum response rate (EF50), while the threshold used by Moreau, Stout and colleagues was based on the observation that the amount of nose pokes per minute in absence of stimulation never exceeded fifteen^32^.

**Table 2.** ICSS methodology of included experiments synthetized by co-authorship network.

| Co-authorship network | Area | Coordinates | Manipulandum | Electrodes | Stimulation | Pulse (ms) | Train (ms) | Trial (s) | Method | Criterion* | Outcome |
| --- | --- | --- | --- | --- | --- | --- | --- | --- | --- | --- | --- |
| **Baker-Bielajew**<br>6 exp/2 studies | VTA | AP 4.8 mm posterior to bregma;<br>ML 0.7 mm;<br>DV 8 mm | Lever press | Monopolar | Cathodal | 0.1 | 300/500 | 60 | Rate-frequency curve shift | Thresholds did not vary by more than 0.1 log<sup>10</sup> units for 3 consecutive test days. | Half maximum (M50) |
| **Lin**<br>1 exp/1 study | MFB<sup>α</sup> | AP -0.5 mm;<br>ML +/- 1.7 mm;<br>DV -8.3 mm | Rotating wheel | Bipolar | Cathodal | 0.1 | 500 | 7.5 | Discrete trial current-threshold | Less than 10% variation in thresholds for 3 consecutive tests | % baseline *Threshold:* Mean of ascending /descending series' thresholds<sup>β</sup> |
| **Moreau-Stout**<br>13 exp/7 studies | VTA | AP 1.5-2 mm from lambda;<br>ML 0.3 mm,<br>DV 8.5 mm | Nose Poke | Bipolar | Alternating | 0.1 | 500 | 120 | Rate-frequency curve shift | Thresholds varied by less than 10 % for 3 consecutive daily test sessions | % baseline *Threshold:* Frequency eliciting 15 nose-pokes per min<sup>γ</sup>. |
| **Nielsen**<br>3 exp/1 study | VTA | AP 3.2 mm anterior to the interaural line;<br>ML 0.6 mm;<br>DV 8.2 mm | Nose Poke | Bipolar | Alternating | 0.1 | 500 | 120 | Rate-frequency curve shift | No increase or decrease trends in maximum response and EF50 varied by less than 10-12% | Mean shift in M50 (log) |

**Abbreviations:** ms, milliseconds; s, seconds; VTA, ventral tegmental area; MFB, middle forebrain bundle; AP, antero-posterior; ML, medio-lateral; DV, dorso-ventral; exp, experiment. Experiments belonging to each group are detailed in *Table 1*.
(\*) Criterion used to indicate the animals learned to self-stimulate.
(α) At the level of the posterior lateral hypothalamus.
(β) Mean intensity calculated from four descending and ascending series of trials (two each), indicating the current at which the animal began to respond positively to the stimulation.
(γ) Based on the observation that without any stimulation, the response level never exceeded 15 nose-pokes per minute.

#### Sucrose preference test

Unfortunately, there was insufficient data to assess whether an increased self-stimulation threshold correlated negatively with sweet consumption. Two studies employed the SPT^39,42^ but only one of them measured the sweet preference in the same group of animals that were assessed for intracranial self-stimulation^39^. In this study by Baker and collaborators, stressed female rats of two common stocks (Sprague Dawley and Long-Evans) underwent 20 hours of food and water deprivation before being offered both water and a 1% (w/v) sucrose solution. After 24 hours, the extent to which the sweetened solution was consumed and preferred over water (the sucrose preference) was measured and reported before and after three weeks of stress. Absolute sucrose intake was reduced but neither the sucrose preference nor the self-stimulation thresholds were altered as a consequence of chronic stress. The authors suggested that the reduced intake reflected an overall reduction in fluid consumption, rather than a specific effect on sucrose intake. This study did not report a correlation between intracranial self-stimulation and sucrose preference. The other study conducted by Nielsen and colleagues assessed the utility of both hedonic measures in different groups of male Wistar and PVG hooded rats exposed to 9 weeks of stress^42^. Sucrose (1% w/v) intake was reduced in stressed PVG rats, although not consistently between weeks, while it remained unaltered in Wistar rats. ICSS thresholds remained unchanged for both rat populations.

#### Risk of bias assessment

All of the included studies suffered from substantial risk of bias, mostly stemming from poor reporting (Figure 3). The average risk of bias score was 3.4 points out of 9 (the lower the number, the higher the risk). All studies reported how the animals were trained in the ICSS procedure and the criteria for when the animals were considered having learned to self-stimulate (ICSS acquisition). Likewise, in all studies but one, it was clear from the reporting that the experimental groups were balanced in key characteristics at baseline. Seven studies reported having confirmed the placement of the electrodes through a histological analysis. Two studies were classified as high risk in this item. In one study, the placement of the electrodes could not be confirmed in some of the animals^42^, while in the other study, it was reported that some of the electrodes were located outside the intended target area^40^. Excluding the studies employing a within-subject design, only two studies out of nine of those using a control vs exposed design were deemed to have a low performance bias because they reported implementing measures to avoid indirectly exposing control animals to stress. Important strategies that minimize bias such as randomized outcome assessment and blinding were not reported in the included studies and only one study mentioned to have randomly assigned their animals to the experimental groups. Similarly, only one study was considered to have a low risk of attrition bias. For the other studies, it was difficult to assess if there were any missing data due to inadequate reporting. Indeed, over half the studies (six) did not report all descriptive statistics (group size, mean, and variation). Only three studies had a low risk of reporting bias. In summary, although the experiments were technically sound, they are limited by unclear risks of bias. Several studies suffered from issues related to poor reporting and there was a lack of clarity on the use of important strategies for minimizing bias.

**Figure 3.**
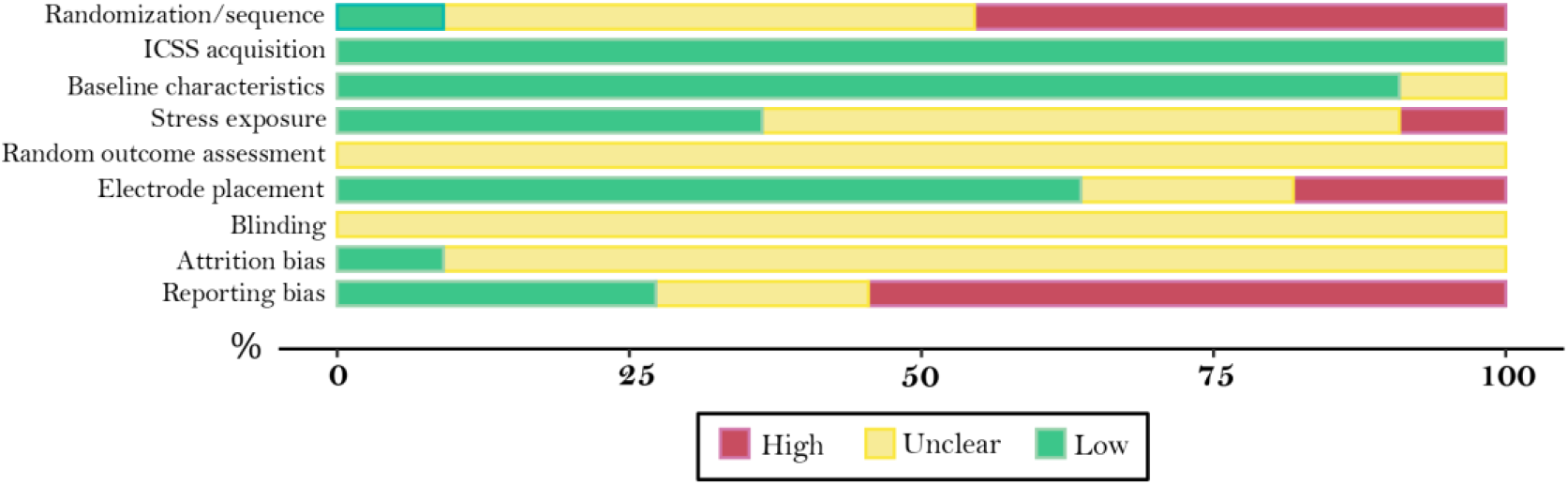
Traffic-light plot of the risks of bias. The risks of bias were assessed using a modified version of the SYRCLE’s risk of bias tool^25^. Each criterion is described in the Supplementary table 4. The colored bars represent the percentage of studies for each item with low (green), unclear (yellow), and high (red) risk of bias.

### Data synthesis

Rats showed a significantly increased self-stimulation threshold after stress when compared to baseline measurements (Supplementary figure 1). When the data was pooled using the two-level meta-analysis, we found that, on average, chronically stressed rats showed a significantly increased self-stimulation threshold compared to unstressed controls (SMD: 0.97; 95% CI: [-0.52, 1.40]; *p* < 0.0001)(Figure 4). Notably, there was significant variation between experiments (I^2^ = 74.3%; 95% CI: [61.4%, 82.9%]; *p* < 0.0001). When accounting for the fact that several experiments were conducted by the same co-authorship network (using a three-level meta-analytical model), the overall effect could not be substantiated. Rats did not show a significantly increased self-stimulation threshold after stress, neither when comparing to their own baseline measurements (Supplementary figure 2) nor in comparison to unstressed controls (SMD: 0.59; 95% CI: [-1.02, 2.19]; *p* = 0.33)(Figure 5). The estimated variance components for the comparisons between stress and controls were τ^2^ = 0.86 at level 3 and τ^2^ < 10^-9^ at level 2. This means that 61.71% of the total variation can be attributed to differences in the results between different networks (I^2^ level 3), while variation in the results obtained in different experiments conducted by the same network was negligible (I^2^ level 2) (Supplementary figure 3). This pattern of variation can be easily observed in the forest plot (Figure 5). Individual experiments performed by the same co-authorship network produced fairly similar effect sizes. However, the direction and magnitude of these effects varied across experiments performed by different networks of authors. Three of the co-authorship networks did not observe significant increases in self-stimulation thresholds in rats exposed to unpredictable stress, while Moreau, Stout and their collaborators consistently reported elevated thresholds.

**Figure 4.**
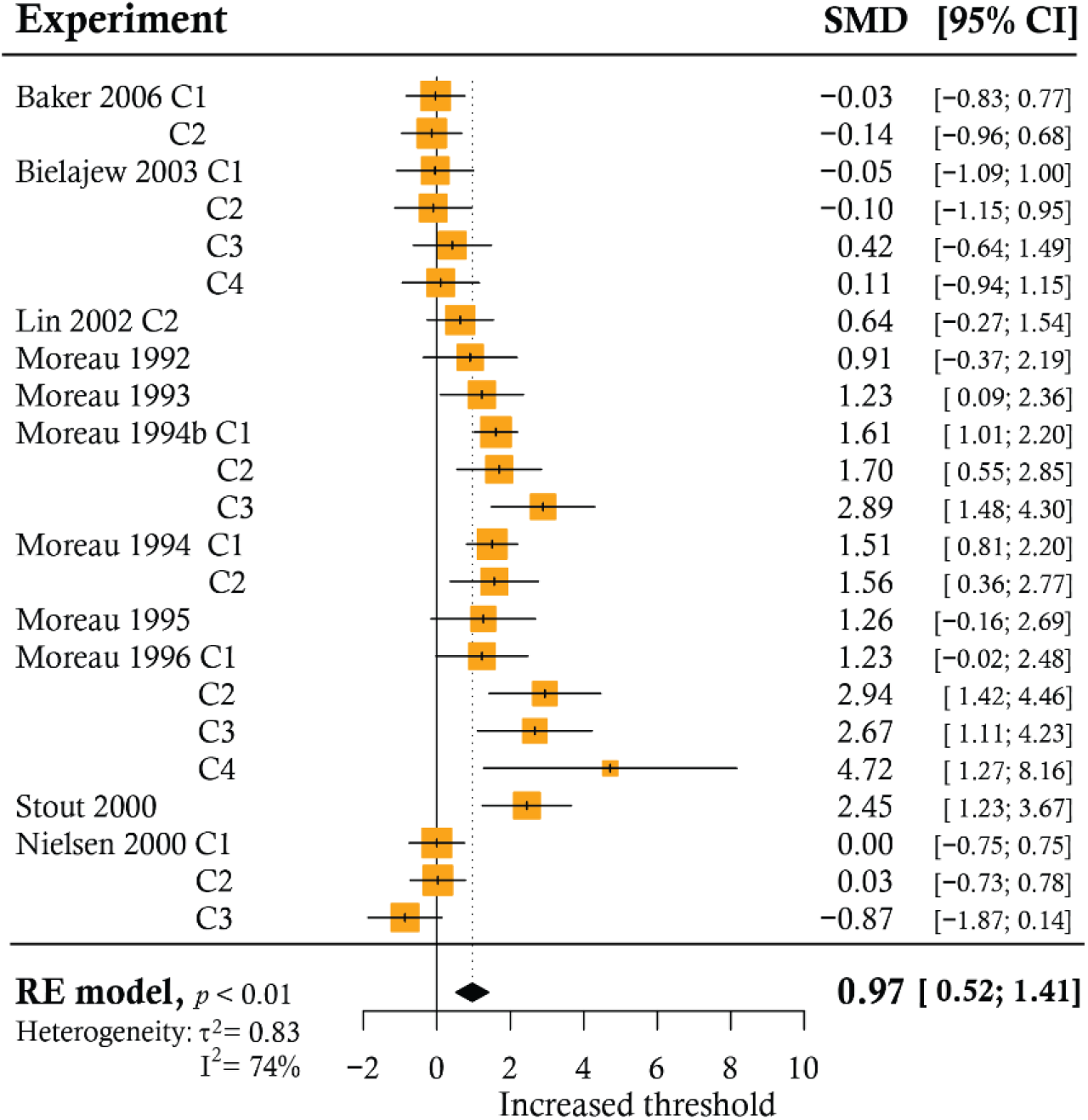
Forest plot of experiments comparing the self-stimulation threshold in stressed animals to unstressed controls. Two-level meta-analysis. The names of the experiments are composed of the main author of the study, the year of publication and an indicator of the number of comparisons (experiments) extracted from a single published study (“Cx”). Each line represents the effect size of an individual experiment with its 95% CI. The sizes of the boxes are proportional to the weight of the experiment in the analysis, which in turn reflects the precision of the study. More precise experiments carry greater weight. The pooled effect and its 95% CI are represented by the black diamond. Studies falling on the right side of the zero line found an increased threshold in stressed animals. Four experiments (Baker 2006 C1, C2; Moreau 1995; Moreau 1996 C4) employed a within-subjects design and pre-stress measurements were used as the control condition. Abbreviations: SMD, standardized mean difference; 95% CI, 95% confidence interval; RE, Random effects.

**Figure 5.**
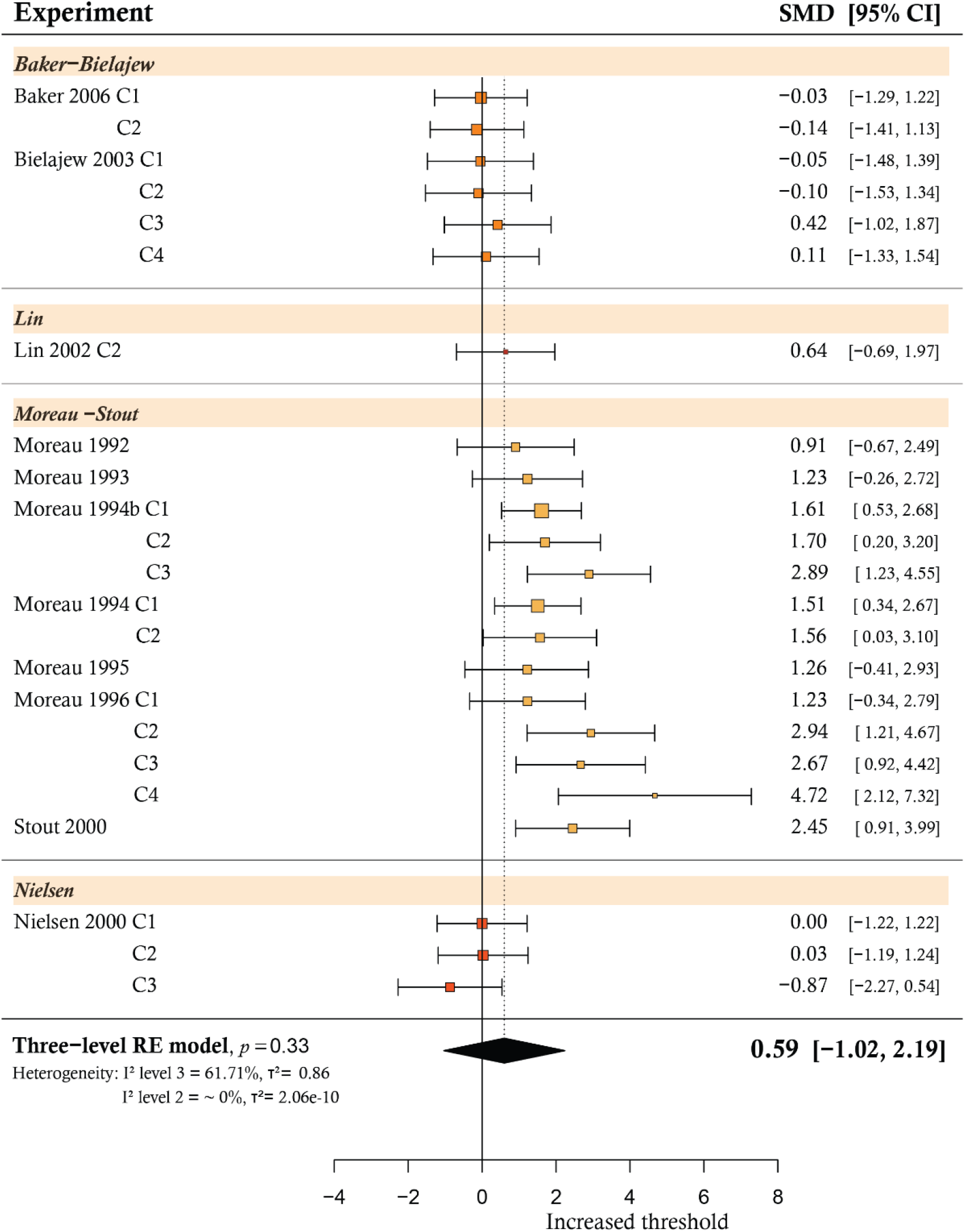
Forest plot of experiments comparing the self-stimulation threshold in stressed animals to unstressed controls. Three-level meta-analysis. The names of the experiments are composed of the main author of the study, the year of publication and an indicator of the number of comparisons (experiments) extracted from a single published study (“Cx”). Each line represents the effect size of an individual experiment with its 95% confidence interval. The sizes of the boxes are proportional to the weight of the experiment in the analysis. The experiments are organized by co-authorship network. The pooled effect and its 95% CI are represented by the black diamond. Studies falling on the right side of the zero line found an increased threshold in stressed animals. Four experiments (Baker 2006 C1, C2; Moreau 1995; Moreau 1996 C4) employed a within-subjects design and pre-stress measurements were used as the control condition. Abbreviations: SMD, standardized mean difference; 95%-CI, 95% confidence interval.

When we compared the two-level meta-analysis to the three-level model, we found that the latter provided a significantly better fit for our data (χ ^2^ = 20.08, *p* < 0.001)(Supplementary table 6). Therefore, we concluded that chronically stressed rats did not show a significantly increased self-stimulation threshold compared to unstressed controls when the individual differences between co-authorship networks were accounted for.

Meta-regressions were performed to assess if the length of stress or the risk of bias score were significant predictors of the magnitude of the effect. Neither the duration of stress (*p* = 0.87) nor the risk of bias (*p* = 0.58) appeared to influence the effect size. (Supplementary table 7). To test the robustness of our results, we conducted two sensitivity analyses. The results remained consistent when we excluded the comparisons where we used pre-stress data as controls and when the highest number within the group size range was used instead of the lowest (Supplementary tables 8 and 9).

No evidence of a small-study effect, or potentially of publication bias, was observed in our data (supplementary material).

### Level of certainty in the body of evidence

We carried out separate assessments for the certainty of evidence for the two-level meta-analysis and the model where between-networks variability was accounted for (**Table 3**). A high level of certainty is assumed at the outset using the GRADE approach. However, since we observed that the included experiments were overall limited by unclear risks of bias, the level of certainty was reduced. The execution of the studies may have introduced systematic errors or inaccuracies that could have significantly affected the results. As a result, for both models, the level of certainty was downgraded to “moderate.” The level of certainty was further downgraded to “low” for the two-level meta-analysis owing to inconsistency of results. The effect sizes of individual experiments were highly heterogeneous. This was not considered an issue for the three-level meta-analysis because this model helped in reducing the impact of heterogeneous results by accounting for an important source of variability in the results: the co-dependency between effect sizes coming from the same network of researchers. No other factors were considered serious enough to merit further downgrading of the certainty of either model.

**Table 3.** GRADE summary of findings table.

| Intracranial self-stimulation after chronic unpredictable stress |  |  |  |  |  |  |
| --- | --- | --- | --- | --- | --- | --- |
| **Population:** Laboratory rats |  |  |  |  |  |  |
| **Intervention/Exposure:** Chronic unpredictable stress |  |  |  |  |  |  |
| **Comparison:** Unstressed controls |  |  |  |  |  |  |
| Outcome | Number controls | Number stressed | Model | SMD (95% CI) | Heterogeneity (%) | Quality of evidence |
| Intracranial self-stimulation | 213.5* | 243.5* | Standard | 0.97 [0.52, 1.41] | 74.3 [61.4, 82.9] | ⊕⊕⊖⊖<br>Low<sup>†</sup> |
|  |  |  | 3-level | 0.59 [-1.02, 2.19] | Level 2<sup>α</sup>: 0<br>Level 3<sup>β</sup>: 61.71 | ⊕⊕⊕⊖<br>Moderate<sup>‡</sup> |
| **GRADE levels of certainty** |  |  |  |  |  |  |
| **High:** We are highly certain that the actual effect is similar to our estimated effect |  |  |  |  |  |  |
| **Moderate:** It is probable that the true effect is similar to our estimate, but there is a chance it could differ significantly |  |  |  |  |  |  |
| **Low:** There is the possibility that the true effect differs markedly from our estimate |  |  |  |  |  |  |
| **Very low:** There is a high probability that the true effect differs significantly from our estimate |  |  |  |  |  |  |
| Foot notes: |  |  |  |  |  |  |
| * In four experiments, the pre-stress measurements were used as the control condition, in these experiments the group size was divided by two to avoid double counting animals. Odd group sizes resulted in fractional numbers. |  |  |  |  |  |  |
| α Within- network heterogeneity. |  |  |  |  |  |  |
| β Between- network heterogeneity. |  |  |  |  |  |  |
| † Downgraded because of risk of bias and inconsistency (see text for further details). |  |  |  |  |  |  |
| ‡ Downgraded because of risk of bias. |  |  |  |  |  |  |

## Discussion

In the present study, we did not find support for the hypothesis that chronic unpredictable stress (CUS) is associated with an increased self-stimulation threshold in rats when accounting for differences between research groups. At a glance, indiscriminately combining all available data, it might appear as if stressed animals increased their ICSS threshold compared to their pre-stress values and unstressed controls. However, these findings could not be substantiated when we added an extra level to the model to account for differences between co-authorship networks. This is an important adjustment in situations like the one we have been investigating, where there is only a small number of studies originating from a limited number of networks of researchers. Some groups of collaborators may find certain outcomes more often than others, either because of the specific methods they employed, the population under study, because of some prior knowledge, or due to the way they conduct, analyze, or report their experiments^29,44^. If one group contributes a disproportionate amount of data to a meta-analysis, the outcome can be distorted^29^. In our study, we observed that one network of authors, composed by Moreau, Stout and colleagues^32–38^, conducted considerably more experiments than the rest, contributing over 50% of the data. What is more, six of their experiments conducted between 1992 and 1996, marked the first evidence of the effect of chronic mild unpredictable stress on self-stimulation in rats^32–37^. These initial studies consistently reported an increased threshold after chronic stress exposure. However, the majority of the publications that followed and that aimed to replicate these findings did not yield the same results. Only one publication was the exception; a study that again originated from the same pioneering co-authorship network^38^.

This finding is crucial as it indicates that ICSS has so far only worked as an outcome ‘in the right hands’. Although, the specific reasons for why this is the case are difficult to ascertain. Regrettably, the limited amount of data prevented us from conducting an extensive analysis to explore potential sources of heterogeneity in the results. While we observed that the duration of stress was not a significant predictor of the observed effects, further insights remain speculative. The only outstanding difference among the studies with positive results, in terms of subject characteristics, was the stock that was used. Moreau-Stout consistently used Wistar RoRo rats in their experiments, a Wistar stock originated in Füllinsdorf, Switzerland from Biological Research Laboratories limited (now Inotiv). These rats have no record of being in use today. However, studies with negative results employed Wistars from other vendors. This begs the question whether this particular Wistar stock was differently susceptible to the stress regimen. The answer to this question remains unclear. Methodologically, no specific factor appears to be uniquely employed in studies reporting positive results. All studies employed a method that has been validated and gives a reliable measure of reward sensitivity that is not confounded by motor performance^17^. Indeed, one of the networks (Nielsen) reporting negative results employed a very similar ICSS protocol to that of Moreau-Stout, differing only in how they determined reward thresholds and the specific structure within the VTA that was targeted. Other studies also differed in the subregion that was stimulated. Since different subregions of the VTA have been shown to be heterogeneous, with neuronal populations differing in their molecular and electrophysiological properties and projections^45^, the influence of the location of the electrodes on the ICSS results cannot be excluded. However, Nielsen and colleagues argue that it is unlikely for this factor to explain why they could not find the same effect as Moreau and colleagues. In a small subset of stress-exposed rats (1-2 per group), stimulation thresholds did progressively increase and there was no apparent difference in where the electrodes were placed compared to the other rats^42^. These findings led Nielsen and colleagues to argue that it is more likely that individual differences in response to stress is what may have contributed to the variability^42^. In the other studies, no single factor (sex, strain, stress protocols, or type of ICSS procedure) was considered to be sufficient to explain why the results by Moreau and colleagues were not replicated^39–41^.

The differences in effects can also not be explained by noticeable differences in the quality of the studies. The risk of bias score was not a significant predictor of the magnitude of the effect and we observed that, in general, all the included studies were technically sound but had major pitfalls in terms of potential bias. There was a considerable proportion of unclear and high risks associated with poor reporting and a lack of clarity on whether key strategies that reduce bias were implemented. This clearly impacts how much trust one can have in the results of the included studies, and, as a consequence, the overall effect estimated by pooling them. This is the main reason why the level of confidence in the estimates is moderate, to acknowledge that there is a chance the true estimated effect could differ significantly^31^. This is not the first time we have seen that the quality of the studies decreases the certainty in the conclusions drawn from preclinical studies in the field^10^ and, sadly, we are not the only ones^46^. In fairness, the studies included in the present review were published years before reporting guidelines, such as ARRIVE^47^, were published. Notwithstanding, we would like to take this opportunity to yet again remind researchers of the importance of a clear and transparent reporting. Informative and transparent reports increase reproducibility and replicability of study findings and will increase the robustness of future systematic reviews and meta-analyses.

The CUS model is old and very popular. A review in 2017 documented more than 1300 publications from over 30 countries, with over 200 publications in the year 2015 alone^6^. This is in agreement with our recent systematic search in 2021 where we found over 1000 publications involving laboratory rats^10^. The model is used under the premise that its effect on how much a sweet solution is preferred over water is a reliable measure of anhedonia. Yet, reports to the contrary have appeared in the literature since the early years of the model^l48–53^, and a recent user survey showed that at least 25% of laboratories have had problems reproducing the CUS-induced effects on the SPT^54^. In fact, our systematic review provided evidence that additional factors unrelated to the likeability of the sweet solution confound its results^10^. However, Paul Willner has reaffirmed over the years that anhedonia is a likely explanation for the CUS-induced changes in the sucrose preference test because this conclusion was not solely based on findings in behavioral tests dependent on consummatory behavior^6,12^. He specifically cites the studies conducted by Moreau-Stout that found an effect of CUS on intracranial self-stimulation. This additional support in the early days of the model, and in the face of reproducibility problems with the SPT, likely aided in the widespread adoption of the model and its popularity to this day^6^. However, as we could not find an overall consistent effect on self-stimulation thresholds from CUS, this argument falters.

Considering that the effect on self-stimulation is regarded as an important piece of evidence supporting the validity of the model, it is intriguing that no further replication attempts were made past 2006. It is fair to assume that the challenges associated with conducting such experiments may have played a role. Intracranial self-stimulation is an invasive, time-consuming and technically demanding method that requires specific equipment and training^17^. This naturally makes it a less appealing choice for evaluating the effects of CUS when non-invasive, less labor-intensive alternatives are available. Even so, and in light of the concerns about the reliability of the SPT^10^, the current lack of explanations as to why CUS has only shown effects on ICSS in the hands of specific researchers under particular conditions does increase the uncertainty regarding the capacity of the model to reliably induce anhedonia.

The second objective of our investigation was to analyze studies that have measured sweet consumption (by means of the SPT) and self-stimulation thresholds in the same group of stress-exposed rats. We hoped to assess whether these two variables were inversely related. In other words, we wanted to see if rats showing a decreased preference for sweetened water also showed an increase in their self-stimulation threshold, two outcomes argued to reflect the anhedonic effects of the model^12^. Unfortunately, only two studies have conducted both tests in rats^39,42^. Only one measured both outcomes in the same cohorts^39^. Regardless of the design, both studies concluded that the SPT was unreliable and that at least sucrose intake seemed to not co-vary with self-stimulating behavior. So Willner’s argument is further undermined by the fact that the few studies that evaluated the effects of CUS through both ICSS and in the SPT could not find a relation between the two methods. It is worth noting that neither of these studies provide evidence that a change in self-stimulation threshold is correlated with a specific threshold of sucrose preference. Despite this, it is not unusual to encounter statements in the literature that claim that rats with preferences below 65% have also shown an increased threshold for ICSS^3^. It would be of interest to assess how reliable other tests of anhedonia (such as the conditioned place preference test, or self-administration of sucrose) have been with respect to the effects of the CUS model. Understanding the extent to which the model induces hedonic changes in other tests of anhedonia will be key for outlining the strengths and limitations of the model as one of stress-induced anhedonia. The CUS model, despite its name, is not a mild procedure and can cause considerable distress to laboratory animals. Thus, employing this model for inconclusive research is unethical. It is crucial to ensure its validity and translational value if we are to avoid causing unnecessary suffering and waste of animal lives for little gain.

### Limitations

This investigation was limited to studies in rats, as we focused on the species the model was originally created for. Data from other species, such as mice, could have provided additional valuable information. The criteria used to define a research group considered only connections easily identified through co-authorships in the included reports. Groups might have been different if lifetime collaborations had been taken into account^29^. However, we found no indication of collaboration between the different researcher networks in the included reports.

### Conclusions

In summary, the sucrose preference test and intracranial self-stimulation are two methods used to assess anhedonia in animal models of depression. The former is commonly used to validate the chronic unpredictable stress model, although its reliability is debatable. Intracranial self-stimulation is an alternative method that has the advantage of directly assessing the sensitivity of the brain’s reward system to an electrical stimulation. An increase in the threshold for intracranial self-stimulation after stress is often cited as evidence that the unpredictable chronic stress model induces anhedonia. However, in our study, we did not find support for this. While the experiments conducted by the pioneering network of collaborators reported a significant effect, attempts at replication by others did not yield the same results. What is more, limited evidence points towards a lack of correlation between sweet consumption and intracranial self-stimulation, although this has scarcely been explored. Unfortunately, the limited number of studies did not allow us to explore possible explanations to the replication failures; no factor was observed to be unique to experiments with positive results. The included experiments were limited by unclear risks of bias, with several studies suffering from issues related to poor reporting. Important strategies that minimize bias, like blinding of the experimenters, were virtually completely absent in the reports of the studies. Uncertainties persist regarding the unpredictable chronic stress model’s consistency at inducing anhedonia in rats.

## Author’s Contributions

Jenny Berrio and Otto Kalliokoski conceived and planned the systematic review. All authors carried out the review, discussed the results and contributed to the final manuscript. Jenny Berrio performed the statistical analyses.

## Acknowledgements

We would like to thank the CAMARADES group for their helpful advice in regard to the 3-level meta-analytical model.

