## Supplementary material for "The threshold for intracranial self-stimulation does not increase in rats exposed to chronic unpredictable stress - a systematic review and meta–analysis"

Jenny P. Berrio, M.D.[1]; Jenny Wilzopolski, DVM.[2]; Katharina Hohlbaum, Ph.D.[2]; Otto Kalliokoski, Ph.D.[1]

#### Methods

##### Registration and open access data

Access to the pre-registered protocol can be found here:
https://www.crd.york.ac.uk/prospero/display_record.php?RecordID=425916

Access to additional files and data sets can be found here: https://osf.io/9cnwj/

Throughout the text you will find references to specific files found in this repository.

##### Deviations from the pre-registered protocol

Our initial analysis plan prepared for a scenario where no control groups were incorporated in the design of the studies and only pre-post outcome data of stressed groups were available for the meta-analysis. However, the use of pre-post SMDs has been advised against given the uncontrolled nature of this design and the risk of erring when estimating effect sizes[1,2]. With this in mind and considering that 19 of the 23 included experiments used a control versus exposed design, we decided to present the results comparing unstressed to stressed animals as the primary analysis. All the extracted data have been made available online (https://osf.io/9cnwj/) and the results of the pre-post analysis are presented in the supplementary material. In order to account for the influence of co-authorship network in our data, a three-level random effects model was added to the analyses and compared to the standard two-level model. One more sensitivity analysis was carried out to assess the reliability of our results when experiments without a control group were omitted. The analyses planned with the outcome of the sucrose preference test were not possible because there was not enough data to conduct them. As the use of SMD is linked to an artificial distortion in the funnel plot[3], we initially intended to employ mean differences when feasible. However, due to varying outcomes across experiments, this approach was not possible. Therefore, the Egger's test with the corrected standard error (SE) proposed by Pustejovsky and Rogers was applied to prevent a false positive asymmetry due to the correlation between the SMD and its standard error[4].

#### Search strategy

In all databases, the search encompassed the time span from 1987 (marking the first Chronic unpredictable stress –CUS- protocol) up until May 2023, without any additional predetermined limits. Each search string was composed of keywords and indexed terms, which were tested and improved to eliminate terms that yielded no results or significantly decreased the accuracy of the search. The final three strings were merged to obtain the list of potential studies in each database.

The complete search strings were as follows:

##### *Pubmed*

(a) "Rats"[Mesh] OR "Rat"[tiab] OR "Rats"[tiab] OR "Rattus norvegicus"[tiab]

(b) "chronic unpredictable mild stress"[tiab] OR "chronic unpredictable stress"[tiab] OR "chronic mild stress"[tiab] OR "mild stress"[tiab] OR "Chronic Unavoidable stress"[tiab] OR "Unpredictable Chronic stress" [tiab] OR “chronic variable stress”[tiab] OR “chronic varied stress”[tiab] OR “chronic variate stress” [tiab] OR “chronic stress depression model”[tiab] OR “ Chronic stress model”[tiab]

(c) Self Stimulation"[Mesh] OR "self-stimulation" [tiab] OR "brain stimulation reward" [tiab] OR "brain reward" [tiab] OR "stimulation reward" [tiab] OR "ICSS" [all] OR "Self-stimulate" [tiab] OR "brain stimulation" [tiab]

##### *Embase*

(a) exp rat/ or "rats".mp. or "rat".mp. or "Rattus norvegicus".mp. or "R. norvegicus".mp.

(b) exp chronic unpredictable stress/ or "CUMS".mp. or "chronic unpredictable mild stress".mp. or "chronic mild stress".mp. or "mild stress".mp. or "Chronic Unavoidable stress".mp. or "chronic unpredictable stress".mp. or "Unpredictable Chronic stress".mp. or "chronic variable stress".mp. or "chronic varied stress".mp. or "chronic variate stress".mp. or "chronic stress depression model".mp. or "Chronic stress model".mp.

(c) self stimulation/ or  self stimulation.mp. or brain depth stimulation/ or reward/ or brain reward.mp. or stimulation reward.mp. or ICSS.mp or brain stimulation.mp

##### *Web of science*

(a) TS=(rat$ OR "Rattus norvegicus" OR "R. norvegicus")

(b) TS=("chronic unpredictable mild stress" OR "chronic unpredictable stress" OR “chronic mild stress” OR “mild stress” OR “chronic exposure to stress” OR "Unpredictable Chronic stress" OR “chronic variable stress OR “chronic varied stress” OR “chronic variate stress” OR “chronic stress depression model” OR “Chronic stress model”)

(c) TS=(“self stimulation” OR "brain stimulation” OR "ICSS" OR “stimulation reward” OR “self-stimulate”)

#### Eligibility criteria

Full text of the studies that passed the title and abstract screening were assessed against specific criteria for inclusion and exclusion. Supplementary table 1 presents the reasons of exclusion during screening.

**Supplementary table 1**. *Reasons for exclusion at each stage of screening.* The order of priority of each criteria is represented by the order they are presented at each stage.

| Title and abstract | Full-text screening |
|---|---|
| **Wrong publication type:**<br>✓ Not an original study<br>✓ Systematic review/Meta-analysis | **Wrong publication type**<br>✓ Not an original study<br>✓ Systematic review/Meta-analysis<br>✓ Not an animal study (in vitro/in silico/clinical study). |
| | **Not easily readable**<br>✓ Article is not easy to comprehend, even with the use of a translation tool |
| **Wrong study design:**<br>✓ Not an animal study | **Wrong population:**<br>✓ Mice, pre-weaned rats, mutant/genetically modified strains |
| **Wrong population:**<br>✓ Not a study on live laboratory rats | **Wrong intervention:**<br>✓ The study does not employ a stress paradigm<br>✓ The study employs other forms of chronic stress different from CUS |
| **Wrong intervention:**<br>✓ The study does not employ a stress paradigm<br>✓ The study employs other forms of chronic stress different from CUS | **Wrong outcome:**<br>✓ Study does not use intracranial self-stimulation as part of the battery to assess anhedonia in the CUS model. |
| **Wrong outcome:**<br>✓ Study does not assess intracranial self-stimulation | **Wrong design:**<br>✓ Study did not use a **"control vs exposed"** design or where the exposed group act as their own control (**within-subject design**)<br>✓ CUS was not administered exclusively for two weeks<br>✓ CUS was not administered exclusively: Studies that at the start of CUS protocol, or while it is being implemented, co- |

| | |
|---|---|
| | administer addictive substances or any other drug, compound, or model-inducing intervention AND do not include exposed groups vehicle-treated.<br>✓ CUS was applied following another chronic stress protocol or model-inducing intervention AND do not include exposed groups in which CUS is exclusively applied. |
| ***Abbreviations:** CUS, Chronic Unpredictable Stress | |

#### Reference list check

One of the reviewers checked the reference lists of the included studies in search of studies that might have been potentially missed with the initial search. The title of each reference was screened and if the title suggested an article of potential interest, its abstract and full-text were reviewed. If the article reported a study in rats that underwent CUS and that were assessed for intracranial self-stimulation (ICSS), said article was to be full-text screened in the same way as those identified through the database search. However, no study of interest was found through this search.

#### Study details and outcome data extraction

**Supplementary table 2.** *Methodological details and outcomes extracted from each unique experimental comparison*.

| **Methodological details** | |
|---|---|
| **Study ID** | ♣ Title<br>♣ Corresponding author and e-mail address<br>♣ Year |
| **Subject characteristics** | ♣ Strain<br>♣ Sex<br>♣ Age (≤9 weeks, ≤12 months, >12 months)*<br>♣ Weight |
| **CUS model characteristics** | ♣ Length (# weeks) |
| **ICSS characteristics** | ♣ Brain site + coordinates<br>♣ Manipulandum (e.g. lever, nose poke, etc.)<br>♣ Electrodes (monopolar, bipolar)<br>♣ Electrical stimulation (anodal, cathodal, alternating polarity)<br>♣ ICSS method (rate-frequency or rate-intensity or rate-duration curve-shift, discrete trial)<br>♣ Criteria for starting testing<br>♣ Pulse duration (miliseconds)<br>♣ Train duration (miliseconds)<br>♣ Trial duration (Seconds) |

| | |
|---|---|
| **SPT characteristics** [£] | ♣ Type of sweet: Sucrose or Saccharine<br>♣ Type of test: One-botte, Two bottle<br>♣ Length (h) of food and water deprivation<br>♣ Timing: light, dark cycle<br>♣ Length of test (h)<br>♣ Concentration of sucrose/saccharine (% w/v) |
| **Outcomes** | |
| **Stimulation threshold** | ♣ Pre and post-stress values of stressed animals<br>♣ Pre and post-stress values of unstressed animals (if reported)<br><br>Measuring units:<br>1. 'Theta-0' (T0)<br>2. 'locus of rise'<br>3. 'half maximum' (M50)<br>4. % change<br>5. Other |
| **Sweet preference** [£] | ♣ Pre and post-stress values of stressed animals<br>♣ Pre and post-stress values of unstressed animals (if reported)<br><br>Measuring units:<br>Sucrose/saccharine preference (in percent), or<br>Sucrose/Saccharine intake calculated as weight or volume of sucrose solution (g or mL), or<br>Ratio of sucrose/saccharine intake to total water intake, or<br>Sucrose/Saccharine intake in relation to the animal's body weight (g/kg), or<br>Other |
| **Correlation** [α] | ♣ Correlation coefficients between stimulation threshold and Sweet preference/consumption. |
| **Abbreviations:** CUS, chronic unpredictable stress; ICSS, intracranial self-stimulation; SPT, sucrose preference test.<br>* In cases where age was not provided, body weight was used to estimate it using predetermined ranges obtained from growth curves reported by various suppliers (OSF: "Preparation/Weight2Age.xlsx").<br>[£] Only extracted if the study performed the sucrose preference test in the same cohort of animals that were assessed for intracranial self-stimulation.<br>[α] Only extracted if the study performed a correlation analysis between the outcomes of interest. | |

Age was defined as follows:

- **≤ 9 weeks:** P21-P63
- **Adult ≤ 12 months:** P64-P336 (10 weeks to 48 weeks)
- **Adult > 12 months:** > 48 weeks

The following limits were used for defining adults:

| Strain/stock | Female | Male |
|---|---|---|
| Wistar | ≤ 300g | ≤ 400g |
| Sprague-Dawley | ≤ 250g | ≤ 350g |
| Long-Evans | ≤ 250g | ≤ 350g |

| Strain/stock | Female | Male |
|---|---|---|
| Lister-Hooded | ≤ 200g | ≤ 300g |

When the age or weight were reported in a range, the upper limit was selected to estimate the developmental stage.

The authors of the studies that reported incomplete information were contacted in hope of having access to the complete dataset. If no response was received within a week, and after exhaustion of other alternatives for contacting the authors or completing the data, the study was excluded from the analysis.

#### Cases of incomplete data

1. One study[5] did not report a measure of variability (SD or SEM). Author could not be contacted and no other means of obtaining data were possible. **Study was excluded.**
2. One study[6] reported two experiments. The group sizes for one of the experiments was not reported. Authors were contacted by e-mail, but no response was received. **Experiment was excluded.**
3. One study[7] did not report the group size of the control groups. They were guesstimated from the degrees of freedom reported and the smallest number estimated was used to have a conservative approach.

.

**Supplementary table 3.** *Outcome extraction considerations regarding specific data reporting cases and study designs.* These considerations and actions were agreed upon by the researchers prior to starting the dual extraction.

| **Data reporting** | | **Study design** | |
|---|---|---|---|
| **Consideration** | **Action** | **Consideration** | **Action** |
| a) n reported in ranges | Lowest value was used in the analysis | a) Studies with repeated measurements of ICSS | The pre-stress threshold reported closest to the beginning of the CUS protocol was used in the analysis.<br><br>The post-stress threshold reported closest to the end of the CUS protocol was used in the analysis. |
| b) SD is not reported | Standard error of the mean (SEM) or confidence intervals were used to calculate it | b) Studies where a secondary intervention was started during the CUS and no placebo/sham group was used | Data was extracted from the last test performed before the start of the new intervention.<br>The length of CUS is the length in weeks before interruption occurred. |
| c) It is unclear what the error bars represent in figures | SEM was assumed (for a conservative estimate) | c) Studies reporting sucrose preference test | Extracted the pre and post-stress sweet preferences/consumptions closest to the time points of the ICSS thresholds |

| | | | | | |
|---|---|---|---|---|---|
| d) | Study reports different ICSS units | Extract all units by priority:<br>1.'Theta-0' (T0)<br>2.'locus of rise'<br>3.'half maximum' (M50)<br>4. % change<br>5. Other | d) | Studies reporting correlation coefficients and reporting n in range | Lowest value was used in the analysis |
| | | | | **Control vs stress designs** | |
| e) | Study reports different SPT units | Extract all units by priority:<br>1. Sweet preference (%)<br>2. Sweet intake/weight (g or mL/kg)<br>3. Sweet intake (g or mL)<br>4. Ratio of sweet intake to total water intake<br>5.Other | e) | Studies with several control and/or stress groups AND where one-to-one control vs stress comparisons are not straightforward | Data were combined to obtain an aggregate measure for one-to one comparison<br><br>**Exception:** whenever groups vary in variables of interest (see data analysis), the data were extracted for each group in reference to a single control or stress group and the number of animals was adjusted in the shared group by the number of comparisons[8]. |
| | Study reports data for "stress resilient" and "stress susceptible" animals separately | Data were combined to obtain an aggregate measure for the stress group | f) | Studies with repeated measurements of ICSS | The pre and post-stress thresholds for controls will be extracted from the same time points as the ones used for the stressed group. |

### Risk of bias assessment

**Supplementary Table 4**. *Risk of Bias checklist, modified from SYRCLE's risk of bias tool for animal studies*

| Bias | Item | Signaling questions |
|---|---|---|
| Selection | **Randomization/Sequence** | Did the investigators describe randomization or a clear strategy to allocate the experimental units? |
| | **ICSS Acquisition** | Does the study describe how the animals were trained to self-stimulate and states the criteria used to define when animals have successfully learned to self – stimulate before starting testing with drugs or other manipulations? |
| | **Baseline Characteristics** | In studies that employ **a control vs exposed design:**<br>Was the distribution of sex, age and stimulation thresholds balanced for the stressed and control groups? (Either matched at the beginning or reported in the results)<br><br>Studies with a **within-subjects design**: default YES |
| Performance | **Stress exposure** | In studies that employ **a control vs exposed** design:<br>"Did the researchers ensure that the control animals were undisturbed except for activities related to their daily care and not inadvertently exposed to stress?" For example, by housing them in separate rooms under similar housing conditions or by taking measures to ensure the controls were not exposed to any stress if house conjointly? |

| | | |
|---|---|---|
| | | In studies with a **within-subjects design**:<br>Were all animals submitted to the same stress protocol?<br><br>If the article does not state that different animals were exposed to different sets of stressors, this is a default YES |
| Detection | **Electrode placement** | Was the correct electrode placement confirmed by histology in all the animals included in the analysis? |
| | **Random outcome assessment** | Did the investigators test the animals in a random order during the outcome assessment?<br><br>For those studies performing the test simultaneously for all subjects, it is a default YES. |
| | **Blinding** | Was blinding of the outcome assessor ensured? |
| Attrition | **Attrition** | If there are any missing data (e.g. mismatched numbers in methods vs analysis), were they adequately explained and dealt with transparently?<br><br>A--> n in methods match n in results--> YES<br>B--> n in methods DOES NOT match n in results<br>--> Is the mismatch explained?--> YES/NO<br>C--> n in results but n is not reported in methods/experimental design --> UNCLEAR<br>D--> n not reported in results--> check stats report--> n easily deduced from the Degrees of Freedom:<br>-> Yes-->n in methods match with n in results--> YES;<br>n in methods DOES NOT match n in results --> Is the mismatch explained?--> YES/NO<br>->No--> UNCLEAR |
| Reporting | **Reporting** | Does the study report the ICSS outcome with descriptive statistics (group sizes, measures of central tendency) and a measure of variability for all groups? |

#### GRADE assessment

The assessment considered the following factors:

1. **Risk of bias** of the included studies,

2. **Inconsistency**: The degree of heterogeneity in effect sizes across experiments. The overlap between CIs, and the direction and magnitude of individual effect sizes. This was serious if there was large variation on effect sizes, no or minimal overlap in the confidence intervals and significant between experiments heterogeneity ($I^2$).

3. **Imprecision**: The precision of the overall effect. This was considered serious if the pooled estimate was based on few animals or if its confidence intervals were judged to be wide.

4. **Indirectness:** The degree to which the pooled evidence directly reflects our PICO and direct conclusions can be drawn regarding our population, intervention and outcome of interest. This was considered serious if the research does not directly compare the population, intervention and outcome of interest.

5. Evidence of **Publication bias.**

Each of these factors demoted the certainty level by one if found serious.

**Supplementary Table 5**. *Levels of certainty of evidence according to the GRADE approach.* The definitions were adapted from “Using systematic reviews in guideline development: the GRADE approach”[9]

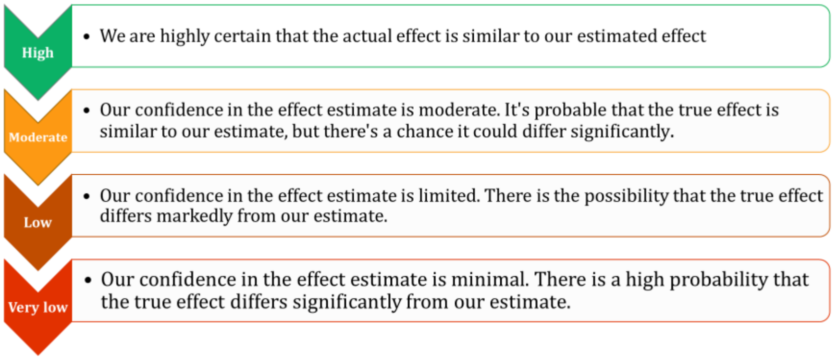


#### Co-authorship networks

Access to the data: *OSF: “Network” & “Nodes”*

The list of authors of the 11 studies that were included was reviewed to know if different publications were authored by the same network of researchers. The first and secondary authors of each paper were extracted and their connections were plotted using cytoscape[10].

#### Data synthesis

Access to the code: OSF: “Code-2SR.qmd”

The following R packages were used in the data analysis and the graphics:

##### Power calculation

Power calculations were performed with the function power.analysis in the R package *dmetar*[11].

A meta-analysis comprising ***four*** studies, each with a sample size of 10, has an 85% power to detect an effect size of 1 standardized effect size (SMD) at an alpha level of 0.05. This calculation was made under the assumption of a random-effects model and high heterogeneity (I2 > 75%). Considering that our previous systematic review of the sucrose preference test found a SMD between the stress and control groups in sucrose consumption/intake of no less than 1 SMD, overall and in multiple comparisons[12], 4 studies were considered to be the minimum requirement for performing an analysis. Likewise, 4 studies per subgroup were deemed necessary to perform subgroup analyses.

##### Data aggregation/synthesis

One study[13] included an experiment where two stress groups were combined using the recommendations in the Cochrane Handbook[8] to obtain a single measure that was compared to the control group.

For the standard analysis of the comparisons between values before and after stress in the stressed group, the group size was split in two to avoid double-counting[8].

The three-level meta-analytical model was performed using the ***rma.mv*** function in the **metafor** package. The calculation of the multilevel $I^2$ was performed with the ***var.comp*** function in **dmetar** package. The graph of the distribution of the total variance was generated by plotting the output of this function. The comparison of the three-level model to one in which the research group level is removed was performed using the **metafor** package. First we employed the ***rma.mv*** function to fit a model in which the level 3 variance, representing the between-research group heterogeneity, is set to zero. The results of this model were then compared to our original model using the ***anova*** function (metafor).

##### Publication bias

The funnel plot and Egger's test for the standard model was performed with the functions ***funnel.meta*** and ***metabias*** in the **meta** package. The argument "Pustejovsky" was selected as the method to test for bias (*method.bias*). The funnel plot for the multilevel meta-analytical model was graphed using the function ***funnel*** in the **metafor** package. The equivalent of the Egger's regression for the three-level model was performed by conducting a three level meta-regression including the corrected standard error of the effect size as a moderator. This modified regression was proposed recently as a good strategy to assess selective reporting (small study effect) while handling dependent effect sizes[14].

#### Results

**Databases:** *(OSF: "Database_R" & "Database_R_main")*

#### Risk of bias

Access to the data:

1. Robvis scale: *(OSF: "RoB_ROBVIS.xls")*
2. RoB score: *(OSF: "RoB_Score.xls")*

#### Data synthesis

##### *Self-stimulation thresholds before and after stress in exposed animals*

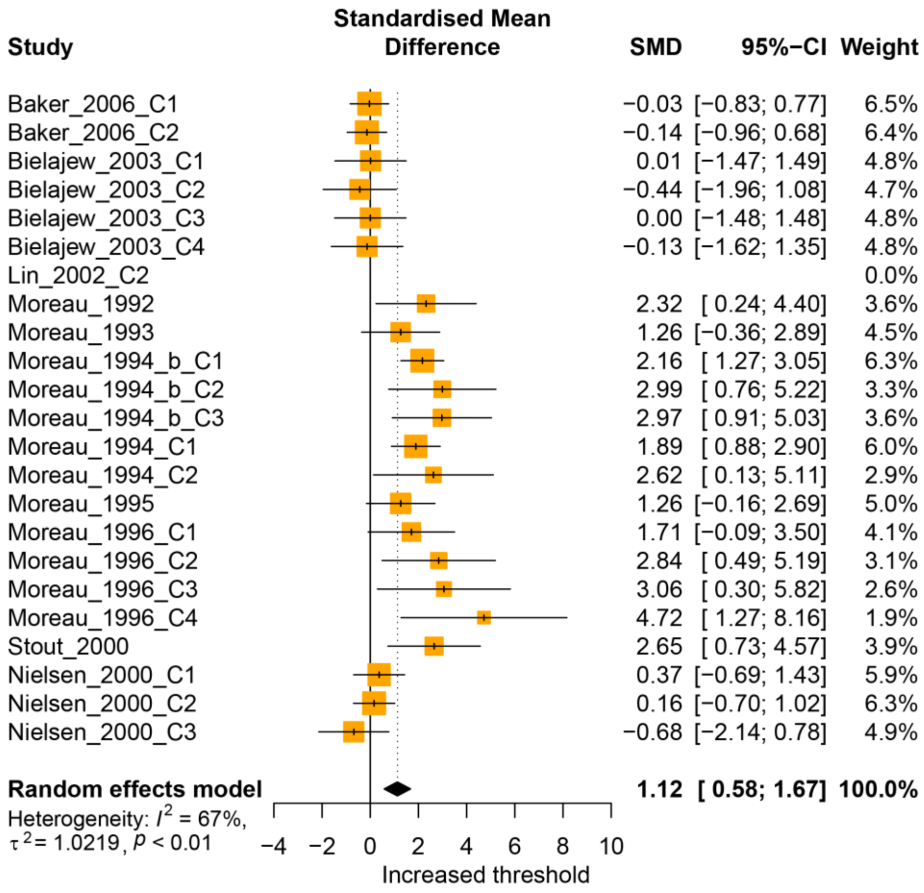


**Supplementary figure 1. Forest plot of 22 experiments comparing the self-stimulation threshold in stressed animals before and after stress. Standard model.** The names of the experiments are composed of the main author of the study, the year of publication and an indicator of the number of experiments extracted from a single published study ("Cx"). Each line represents the effect size of an individual experiment with its 95% confidence interval. The size of the boxes are proportional to the weight of the experiment in the analysis. The pooled effect and its 95% CI is represented by the black diamond. Studies falling on the right side of the zero line found an increased threshold in stressed animals. Total animals: 133 rats. Abbreviations: SMD, standardized mean difference; 95%-CI, 95% confidence interval.

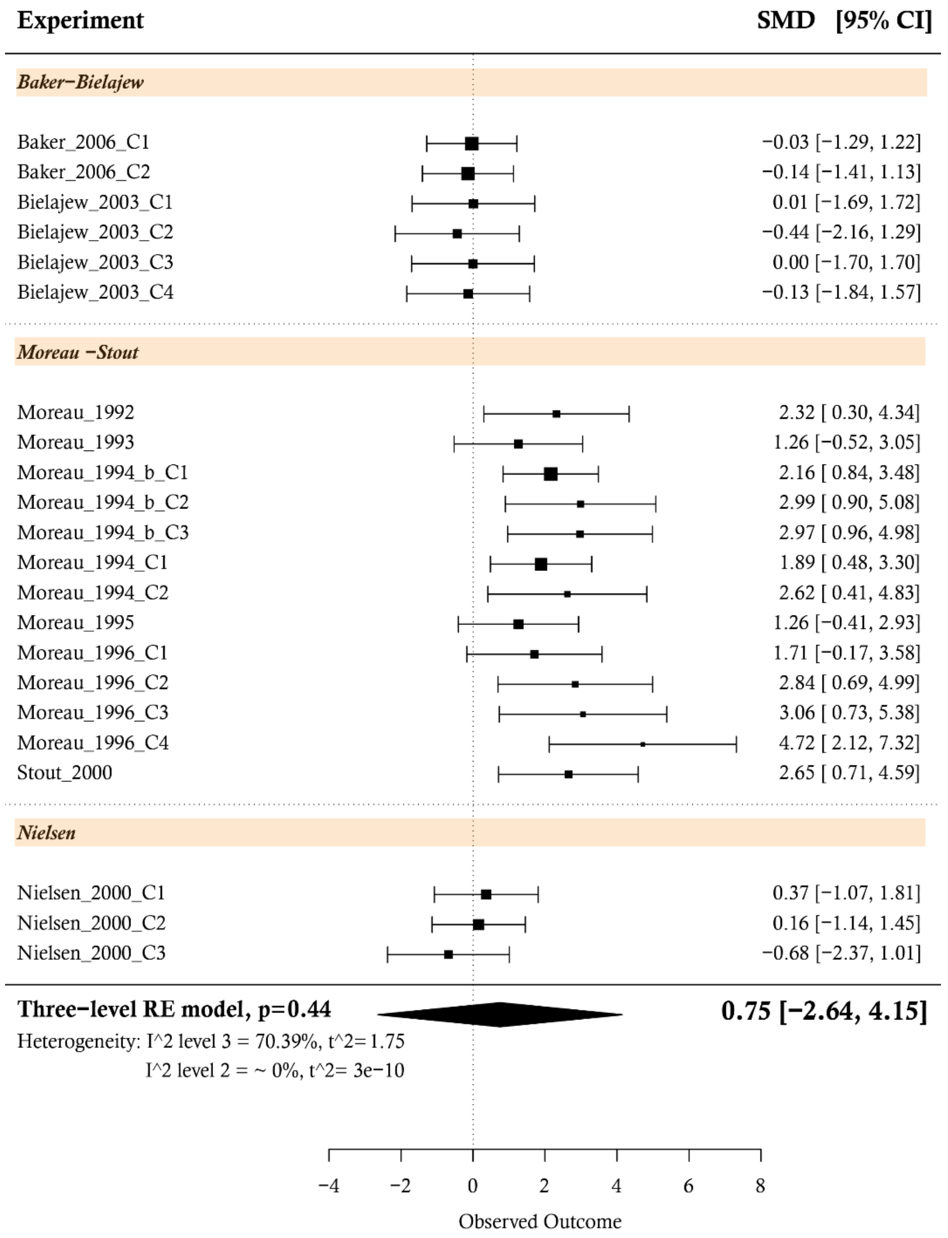


**Supplementary figure 2. Forest plot of 22 experiments comparing the self-stimulation threshold in stressed animals before and after stress. Three-level meta-analytical model.** The names of the experiments are composed of the main author of the study, the year of publication and an indicator of the number of experiments extracted from a single published study ("Cx"). Each line represents the effect size of an individual experiment with its 95% confidence interval. The size of the boxes are proportional to the weight of the experiment in the analysis. The effect sizes are ordered by research group. The pooled effect and its 95% CI is represented by the black diamond. Studies falling on the right side of the zero line found an increased threshold in stressed animals. Total animals: 133 rats. Abbreviations: SMD, standardized mean difference; 95%-CI, 95% confidence interval.

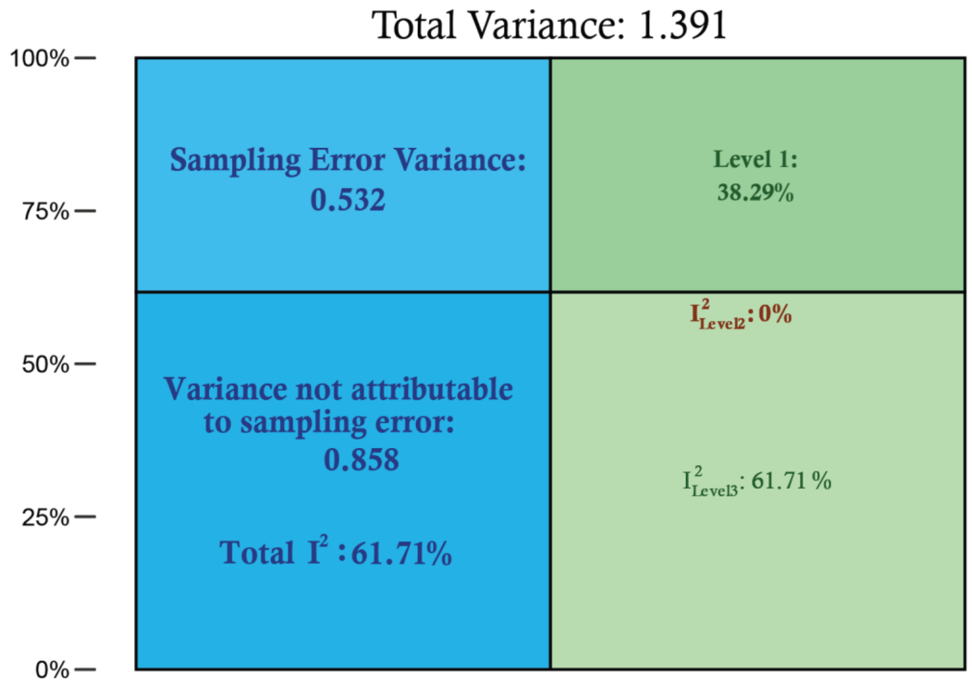


**Supplementary figure 3.** *Plot of the distribution of the total variance in the three-level meta-analysis comparing stressed vs control groups*. The percentage of total variance attributable to each of the three levels is presented on the right side. I2 Level 2 represents the amount of heterogeneity within clusters (research group), while I2 Level 3 represents the between-cluster heterogeneity. The total of the variance in the model that is not attributable to sampling error (61.71%) comes from differences in results across research groups. Percentage of variance within experiments of the same group are ~ 0%.

**Supplementary Table 6**. *Model fit statistics and the likelihood ratio test comparing the three-level with a standard model (reduced).*

| | df | AIC | BIC | LRT | pval |
|---|---|---|---|---|---|
| | | | **Stress (pre–post comparison)** | | |
| **Three-level model** | 3 | 57.09 | 60.23 | | |
| **Reduced model** | 2 | 78.77 | 80.86 | 23.68 | <.0001 |
| | | | **Stress Vs control** | | |
| **Three-level model** | 3 | 56.96 | 60.24 | | |
| **Reduced model** | 2 | 75.04 | 77.22 | 20.08 | <.0001 |

Abbreviations: df, degrees of freedom; AIC, Akaike information criterion; BIC, Bayesian information criterion; LRT, likelihood ratio test; pval, p value.

Foot notes:

The AIC and BIC are two criteria used for model selection, they measure how well a model fits the data. Lower AIC and the BIC indicate better performance. The AIC estimates the model's capacity to predict future values. A lower AIC score indicates that the model can make predictions with nearly the same level of accuracy while needing less information[15,16]. The BIC balances the accuracy of the predictions with the complexity of the model. A lower BIC indicates the model balances accuracy and simplicity[17].

##### Other sources of variability

**Supplementary table 7.** *Summary univariate meta-regression analysis.*

| Variable | Coefficient (SMD) | 95% CI | *t* | p value |
|---|---|---|---|---|
| Length of stress (weeks) | -0.0206 | [-0.27, 0.22] | -0.17 | 0.87 |
| Risk of bias score | -0.1082 | [-0.51, 0.29] | -0.56 | 0.58 |
| Abbreviations: SMD, Standardized mean difference; CI, confidence interval | | | | |

#### Sensitivity analysis

##### Removing pre-post stress comparisons

To test the robustness of our results, we re-ran the primary analyses after removing the experiments were the threshold obtained before stress was used as control (4 experiments removed). Results were similar to those obtained with the main analyses (**Supplementary table 8**). The three-level model was still a better fit for our data ($\chi_1^2$= 15.15, $p$< 0.001)(**Supplementary table 9**).

##### Sample size

Data: *(OSF: "Database_R_S.xlsx")*

When the sample sizes of the experiments were reported in ranges (1 experiment), the lowest value was used for the primary analyses. The meta-analyses were re-run using the largest possible sample size. Results were similar to those obtained with the main analysis. (**Supplementary table 8**).

**Supplementary Table 8**. *Model statistics comparing the primary analysis with the results of the two Sensitivity analysis.*

| Analysis | SMD | 95% CI | *$I^2$ level 2* | *$I^2$ level 3* | p value |
|---|---|---|---|---|---|
| **Standard random effects model** | | | | | |
| Primary | 0.97 | [ 0.52; 1.40] | 74.3% | | < 0.0001 |
| Pre-pos removed | 1.02 | [ 0.54; 1.50] | 74.3% | | < 0.0001 |
| Highest sample size | 0.96 | [ 0.52; 1.41] | 75.1% | | < 0.0001 |
| **Three-level random effects model** | | | | | |
| Primary | 0.59 | [-1.02, 2.19] | 0% | 61.71% | 0.33 |
| Pre-pos removed | 0.60 | [-0.96, 2.16] | 0% | 60.31% | 0.31 |
| Highest sample size | 0.58 | [-1.03, 2.19] | 0% | 62.16% | 0.33 |
| Abbreviations: SMD, Standardized mean difference; CI, confidence interval | | | | | |

**Supplementary Table 9**. *Model fit statistics and the likelihood ratio test comparing the three-level with a standard model (reduced).* ***Sensitivity analysis****.*

| | df | AIC | BIC | LRT | pval |
|---|---|---|---|---|---|
| **Three-level model** | 3 | 45.23 | 47.90 | | |
| **Reduced model** | 2 | 58.38 | 60.16 | 15.15 | <.0001 |

Abbreviations: df, degrees of freedom; AIC, Akaike information criterion; BIC, Bayesian information criterion; LRT, likelihood ratio test, pval, p value.

#### Publication bias - small study effect

The funnel plot visually represents the relationship between precision (standard error) and effect size (SMD or MD) in experiments. Experiments with higher precision should cluster closely around the overall effect. As precision decreases, experiments scatter more widely, forming the characteristic funnel shape. Small studies, historically less likely to be published, can introduce bias, causing asymmetry in the funnel plot shape, known as the 'small study effect´. MD is preferred over SMD because its use is not associated with any artificial asymmetry in the funnel plot[3]. However, in our study, this measure of effect could not be used given the varied outcomes reported in the included experiments.

The influence of small studies[18] was initially assessed in the standard model by visual inspection of funnel plot of the SMD in relation to its standard error (SE) followed by Egger's test with the corrected SE proposed by Pustejovsky and Rogers[4]. The overall pattern looked slightly asymmetrical (**Supplementary figure 3A**). Some of the experiments with higher positive effect sizes (bottom-right corner) are not balanced by similarly sized experiments reporting null effects, or effects in the opposite direction (bottom-left corner), producing a slightly lopsided funnel. Egger's regression test with the Pustejovsky correction did not confirm asymmetry (intercept: -0.99, t: -0.48, p = 0.64). As a *post hoc* analysis, we assessed the small study effect in the three level model. The slightly asymmetrical pattern of the funnel plot was observed again (**Supplementary figure 3B**). Similar to the standard model, the modified regression test did not confirm asymmetry (intercept: -0.93, t: -0.55, p = 0.63).

Assessing the presence of small study effect in the context of the present study is challenging for several reasons. First, most animal studies, contrary to clinical studies, are small studies per se. Group sizes in our included experiments were on average between 9 (control groups) and 12 animals (stressed groups). Second, the impossibility of using an effect measure (MD) that is not associated with artificial asymmetry in the funnel plot brings uncertainty to the interpretation of any observed asymmetry. While using a corrected measure of precision is proposed to overcome this issue when the asymmetry is tested by the egger's test[4], once this corrected version of the standard error is used in the plot, the effect sizes hardly arrange in a funnel plot shape (**Supplementary figure 4).** Third, the experiments that report a positive result (figure 3, orange dots) come from the same research group, while no other group reported significant effects. It is impossible to know whether this group always obtained positive results, or failed attempts were simply not published.

Since no outright asymmetry was observed visually, nor confirmed statistically, we concluded that no small study effect was confirmed. We also considered this to not be extremely relevant in the discussion of our results, since any potential bias in the estimation of the effect that may have been introduced at the research group level was accommodated with in our multi-level model.

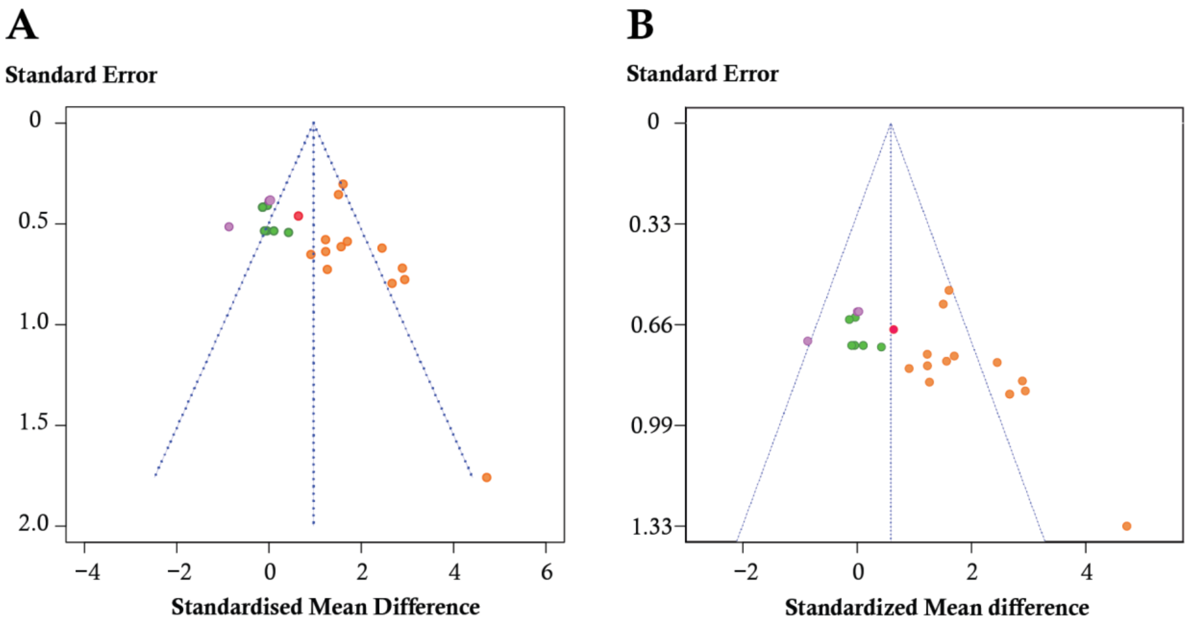


**Supplementary figure 4.** *Funnel plots of experiments when the standard random effects model (**A**) or the multilevel model (**B**) were used.* Experiments compare stressed rats to unstressed controls in their self-stimulation threshold. The observed effect sizes (x-axis) is plotted against their standard error (y-axis). The y-axis is inverted with studies with the lowest standard errors on top. The color of the experiments is given by the research group: orange, Moreau –Stout; red, Lin; green, Baker-Bielajew; and purple, Nielsen. These plots do not use the corrected standard error proposed by Pustejovsky and Rogers.

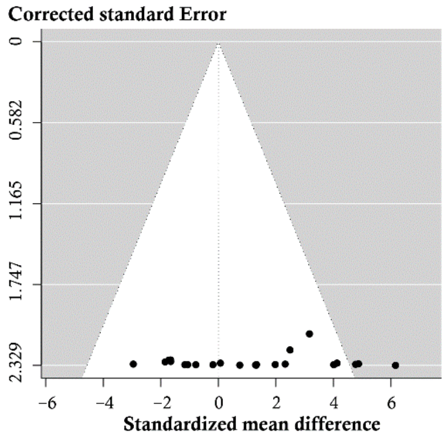

**Supplementary figure 5.** *Funnel plot of experiments when the multilevel model was used and the standard error was corrected according to Pustejovsky and Rogers[15].* Experiments compare stressed rats to unstressed controls in their self-stimulation threshold. The observed effect sizes (x-axis) is plotted against their corrected standard error (y-axis).

#### GRADE assessment

The considerations regarding risk of bias and inconsistency were discussed in the main text.

In our study, imprecision was not judged serious for the standard model because the pooled effect did not cross the no effect line and the confidence intervals were not considered to be wide compared to the ranges of effect sizes found in systematic reviews of animals studies. This same judgement was applied to the three-level model, where we considered the estimate of no effect to be precise. The pooled evidence was a direct reflection of our PICO and the type of ICSS procedures employed to obtain and define reward thresholds are extensively used and known to be reward-selective[19–21]. This is way indirectness was not considered an issue in our investigation. Likewise, the lack of a strong evidence of publication bias refrained us from decreasing our certainty in our results based on it.
